# EGR1 drives sex-differences in glioblastoma tumorigenicity

**DOI:** 10.1101/2025.05.16.654616

**Authors:** Tamara J. Abou-Antoun, Lorida Llaci, Jason P. Wong, Lihua Yang, Nicole M. Warrington, Lloyd Tripp, Jingqin Luo, Brian Muegge, Joshua B. Rubin, Robi D. Mitra

**Affiliations:** Pediatric Hematology-Oncology, Washington University School of Medicine, St. Louis, MO, 63110, USA; Department of Genetics, Washington University School of Medicine, St. Louis, MO, 63110, USA; Center for Genome Sciences and Systems Biology, Washington University School of Medicine, St. Louis, MO, 63110, USA; Department of Pediatrics, Washington University School of Medicine, St. Louis, MO, 63110, USA; Division of Public Health Sciences, Department of Surgery, Washington University School of Medicine, St. Louis, MO, 63110, USA; Biostatistics and Qualitative Research Shared Resources, Siteman Cancer Center, Washington University School of Medicine, St. Louis, MO, 63110, USA; Department of Medicine, VA Medical Center, 915 North Grand Blvd, St. Louis, MO, USA; Department of Medicine, Washington University School of Medicine, Saint Louis, MO 63110, USA; Siteman Cancer Center, Washington University School of Medicine, St. Louis, MO, 63110, USA; Department of Neuroscience, Washington University School of Medicine, St. Louis, MO, 63110, USA

## Abstract

Although progress has been made in treating glioblastoma (GBM), with fewer than 5% of patients surviving more than 5 years after diagnosis. For reasons that are not well understood, females are roughly 60% as likely as males to develop GBM, and female patients consistently respond better to treatment than do males. Understanding the molecular etiology of these sex differences in tumor progression and resiliency to treatment could reveal potent new therapeutic targets, ultimately improving survival of both male and female GBM patients. Here we show that the transcription factor Egr1 is a primary mediator of sex differences in multiple GBM tumorigenic phenotypes. In multivariate analysis, high levels of *EGR1* expression are correlated with shortened survival for male GBM patients only. To investigate the molecular mechanisms underlying this sex difference, we performed a genomic analysis in our established *ex vivo* murine model of sex differences, which showed that the transcription factors Egr1 and Klf5 preferentially recruit the transcriptional activator Brd4 to enhancers in male cells relative to female cells, explaining a previously made observation that Brd4 inhibitors reverse sex differences in GBM. Next, using murine and human primary GBM cells, we demonstrated that the small molecule compound SR18662, which downregulates Egr1 and its downstream target Klf5, abrogates GBM growth, migration, invasion, clonogenicity, and response to radiation in a sex-biased fashion. Finally, we knocked down Egr1 and Klf5 via CRISPRi in both untreated and SR18662-treated GBM cells to reveal the sex-biased anti-tumorigenic effects of SR18662 were largely due to Egr1 downregulation, independent of Klf5 downregulation. This result was replicated in vivo. Our results strongly indicate that an Egr1 regulon is a key determinant of sex differences in GBM. As EGR1 is implicated in the cancer biology of many cancers that also display sex differences in incidence and treatment response, our results are likely to be broadly applicable in oncology.

## Introduction

Glioblastoma (GBM) is the most common lethal primary brain tumor, with fewer than 5% of patients surviving more than 5 years after diagnosis. Additionally, male patients with GBM have a higher incidence^1^ and worse outcome to treatment compared to female patients^2^. This sex difference extends across species, as there is also a male bias in spontaneous GBM incidence observed in dogs^3^. Together, these observations suggest that sex has a conserved and fundamental influence on GBM risk and response to treatment.

The study of sex as a biological variable in basic and clinical research has only recently begun to be appreciated. Investigation into the etiology of sex differences in GBM is likely to reveal novel insights into the incidence, progression, and resiliency to treatment of this disease. Given the effect size of sex differences in GBM, such an approach may also uncover potent therapeutic targets, ultimately improving survival of both male and female patients with GBM. Finally, this approach could also lead to sex-informed treatments, which through greater precision, have the potential to be more efficacious and less toxic than those that ignore sex.

What is the molecular basis of these sex differences in GBM? Sex differences in disease are often ascribed to acute sex hormone actions, but this is unlikely to be the major determinant in GBM because sex differences in brain tumor rates are evident in pre-pubescent, young adult, and post-menopausal adult patients. This, however, does not exclude a potential role for in utero sex hormone epigenetic effects or modulating roles of circulating sex hormones on patterning cancer biology^4^. They may, however, still influence disease phenotype through effects on immunity, metabolism, and aging^5^. Nonetheless, since GBM sex differences are predominantly due to cell-autonomous mechanisms, they can be productively modeled ex vivo. As such, we developed an isogenic *ex vivo* model of GBM in which murine neocortical post-natal day 1 (p1) astrocytes with a combined loss of neurofibromin (NF1) and p53 function. Using this model, we found that sex differences in GBM cancer hallmark pathways are primarily due to cell intrinsic mechanisms^6,7^. This finding has since been further validated in a second murine model that uses in utero electroporation to disrupt function of Nf1 and p53^8^ and in multiple analyses of GBM patient data and primary cells. Since both murine models recapitulate the transcriptional sex differences observed between sexes in human GBM tumors^6^ as well as the phenotypic differences that underlie the greater incidence and shorter survival observed in males^9,10^, these results suggest sex differences in GBM are primarily cell intrinsic. We have since identified a number of molecular pathways that make contributions to these cell-intrinsic sex differences in GBM^11,12^, but we recently achieved a breakthrough when we identified an epigenetic program that, when disrupted by Brd4 knockdown or inhibition, reverses sex differences for nearly all tumorigenic and resiliency phenotypes^13^. Because Brd4 is a general transcription factor that does not contact DNA, it was heretofore unclear which sequence-specific transcription factors (TFs) recruit Brd4^14^, knowledge of which would reveal the pathways directly responsible for sex differences in GBM.

Here, we performed a computational analysis of Brd4 binding data in male and female GBM cells to reveal that the transcriptional regulators Early growth response 1 (Egr1) and Krüppel-like factor 5 (Klf5) recruit Brd4 to male-biased stretch and regular enhancers, respectively. Egr1 is a TF in the EGR family that can get activated by growth and inflammatory factors, ionizing radiation and reactive oxygen species^15^. Interestingly, its role is highly context dependent, as it plays pro-tumorigenic roles in pancreatic cancer^16^ and in hepatocellular carcinoma (HCC)^17^ and anti-tumorigenic roles in rhabdomyosarcoma^18^ and in non-small cell lung cancer (NSCLC)^19^. Its target, Klf5 is a member of the KLF family, and also plays dual roles in cancer. In particular, it plays pro-tumorigenic roles in pancreatic cancer^20^, and in HCC^21^, similarly to Egr1. In other instances, Klf5 inhibits cancer-promoting phenotypes, such as in liver cancer^22^ and in epithelial cells^23^. These contradictory functions of Egr1 and Klf5 in cancers that display sex differences^24–27^ in incidence or outcome suggests that sex needs to be considered when studying their role in cancer.

To study their role in sex differences in GBM, we used both our male and female isogenic murine model as well as human primary GBM cells. We demonstrated that SR18662, a small molecule compound that downregulates both Egr1 and Klf5, inhibits GBM growth, migration, invasion, clonogenic cell frequency, and response to radiation in a highly sex-biased fashion. We knocked down Egr1 and Klf5 using CRISPRi in untreated and SR18662-treated GBM cells to reveal the sex-biased anti-tumorigenic effects of SR18662 were primarily due to Egr1 downregulation. We further established stable knockdowns of Egr1 and Klf5 and replicated these results *in vivo*. Together, these results identify Egr1 as a primary mediator of sex differences in GBM. As EGR1 is implicated in the cancer biology of many cancers that also display sex differences in incidence and treatment response, our results are likely to be broadly applicable in oncology.

## Results

### Egr1 and Klf5 binding motifs are strongly enriched in male-biased stretch or regular enhancers, respectively

Brd4-bound enhancers have previously been shown to play critical roles in maintaining sex differences in GBM^13^. Since Brd4 is a general transcription factor, it must be recruited to enhancers by sequence-specific transcription factors^14^. We therefore reasoned that by identifying the transcription factors that recruit Brd4 to those enhancers associated with sex-biased gene expression, we could find the regulatory pathways directly involved with sex differences in clonogenicity, invasion and metastasis in GBM. Analyses of Brd4-bound stretch-enhancers as previously described^28^ as well as of enhancers^29^, revealed two candidate TFs, Egr1 and Klf5 (**Fig 1A, left panel**), respectively, for further analyses. We focused on these two TFs because of four reasons; 1) Their motifs were both strongly enriched (greater than 3-fold change with a p-value of 1E-7 for Egr1, and >1-fold change with a p-value 1E-5 for Klf5) (**Fig 1A, right panel**) in male Brd4-bound stretch and regular enhancers relative to females; 2) EGR1 expression levels stratify male but not female GBM patient survival (**Fig 1B**); 3) Egr1 is known to directly regulate Klf5, so these two TFs act in the same regulatory pathway and are logically studied together; 4) both Egr1 and Klf5 are known to be critical regulators of biology and response in multiple cancers that also exhibit sex differences in incidence and survival through controlling proliferation and cellular resiliency to stress, areas where we have previously observed major sex differences^5^. Finally, Egr1 and Klf5 can be targeted with the drug SR18662. In conclusion, we hypothesized that inhibiting Egr1 and Klf5 would lead to a decrease in tumorigenic phenotypes in male transformed astrocytes, while no changes would be observed in female transformed astrocytes.

**Figure 1:**
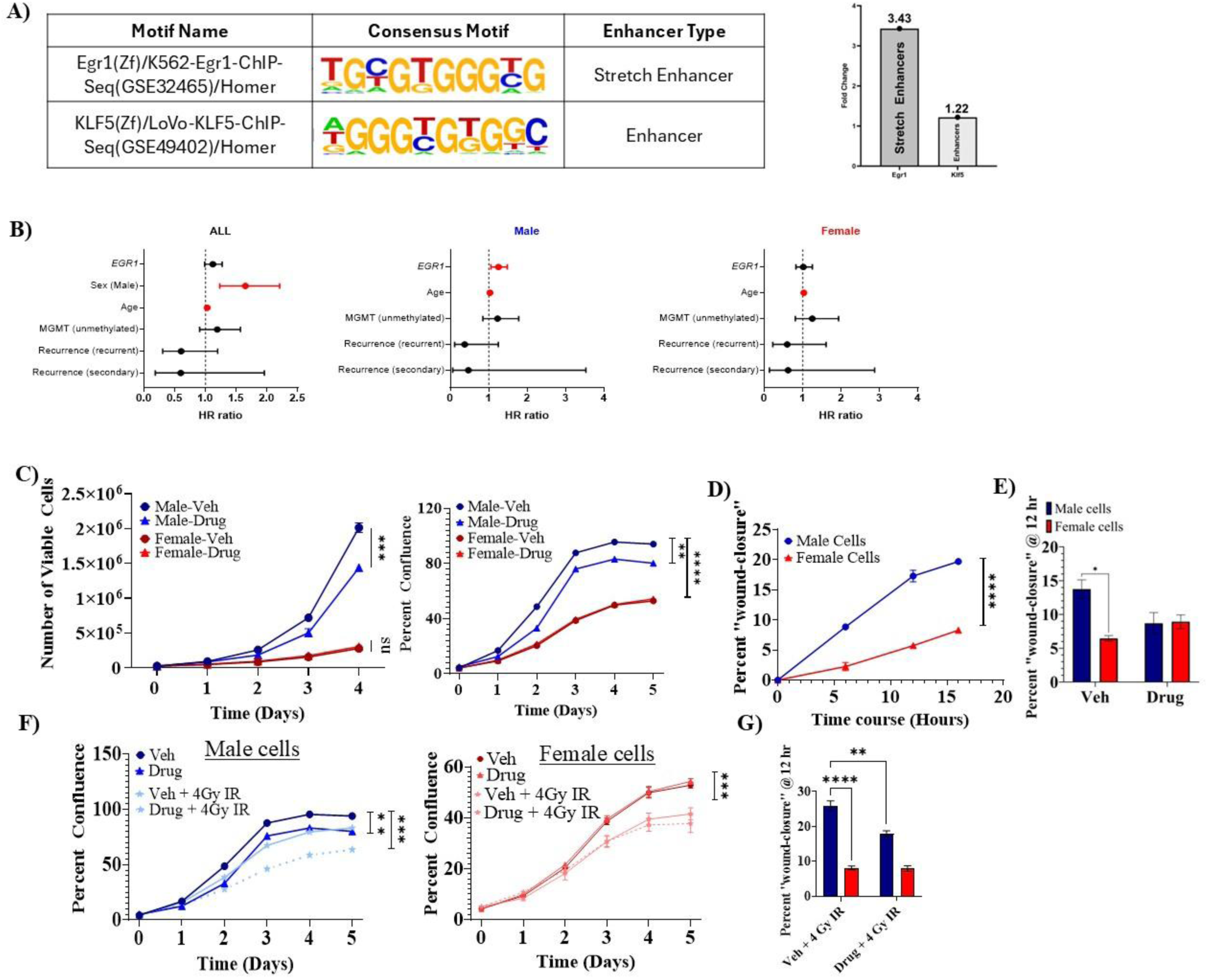
SR18662 treatment induces a sex-biased response on GBM cell growth, migration and irradiation response. **A**) **Left panel**: Top motif enrichment results of Brd4-bound stretch enhancers and enhancers showed Egr1 and Klf5, respectively. **Right panel:** Egr1 motif in Brd4-bound stretch enhancers was 3.43-fold enriched (p value of 1E-7) and Klf5 motif in enhancers was 1.22-fold enriched (p-value 1E-5) in male relative to female respective enhancers**. B)** Forest plot of hazard rations with 95% confidence interval from a multivariate Cox proportional hazard model with the following covariates (*EGR1*, sex, age, MGMT methylation status, and recurrence) on TCGA GBM cohort data from glioVis. **C)** Number of viable male and female GBM cells (left panel) and measure of precent confluence (right panel) after 2µM SR18662 treatment over time. Two-way, repeated measures ANOVA: drug: *p < 0.05* males only; sex: *p < 0.0001*; sex/drug interaction: *p < 0.0001*. **D)** Measure of cell migration as assessed using the “wound-healing” assay to assess percent “wound-closure” in murine male and female GBM cells at 6, 12, and 16 hr post “wound-induction”. Two-way ANOVA: sex: *p<0.0001*; time: *p < 0.0001*; sex/time interaction: *p < 0.0001*. **E)** Measure of cell migration 12 hr post drug treatment. Two-way ANOVA: Sex: *p < 0.05*; Drug: *p = 0.3;* Sex/Drug interaction: *p < 0.05.* F) Time-dependent measure of cell confluency with Drug treatment (2µM SR18662) ± IR (4Gy) over 5 days in murine male (left panel) and female (right panel) GBM cells. Three-way, repeated measures ANOVA: IR: *p < 0.001*; drug: *p < 0.01* (males) *p > 0.999* (females); time: *p < 0.0001*. **G)** Measure of cell migration 12 hr post 4Gy IR ± Drug. Two-way ANOVA: sex: *p = 0.003*; Sex/Drug interaction: *P = 0.004*; IR ± Drug: *p < 0.0001* (Males only). All experiments were conducted in triplicates and repeated at least twice. Error bars represent the mean of all biological repeats ± standard error of the mean (SEM).

### The Egr1/Klf5 inhibitor SR18662 has a sex-biased effect on transformed astrocyte growth, migration, and response to radiation

We next sought to determine if the combined action of Egr1 and Klf5 contributed to sex differences in the tumorigenic phenotype of our isogenic murine model of GBM, which utilizes NF1-/-; DNp53 transformed postnatal day 1 mouse astrocytes. We treated male and female cells with the small molecule inhibitor SR18662, which has been shown to decrease both Egr1 and Klf5 protein expression levels^30^. We performed live cell imaging of low-density cell cultures to quantify the interplay of SR18662 treatment and cell sex on cell proliferation. As illustrated in **Fig. 1C**, males had a significantly higher number of viable cells over time that was significantly higher compared to female transformed astrocytes, underscoring the pronounced proliferative advantage inherent to male cells. Treatment with 2µM SR18662 yielded a sex-biased effect on cell growth; male cells exhibited a significant reduction (28% decline, light blue line in **Fig. 1C**) in growth rate, but female cells were unaffected (**Fig. 1C**, maroon line). This was the smallest dose that resulted in a decrease in proliferation in male cells, while there was no effect on female cells. We also measured GBM cell confluency over time to evaluate the effects of SR18662 over a range of cell densities. We observed a similar trend, where male cells exhibited a significant reduction in both the growth rate and the cell density at which growth saturation was reached after drug treatment, whereas female cells were again unaffected **(Fig. 1C, right panel)**. Importantly, the inhibitory effect of SR18622 on male cell proliferation was dose dependent (**Supplemental Fig. 1A),** providing additional support that downregulation of Egr1/Klf5 is a causal factor in this sex biased phenotype Together, these data suggest Egr1/Klf5 function is required to maintain sex differences in cell proliferation in this DN astrocyte model.

We next quantified the effect of Egr1/Klf5 downregulation on migration, a cellular function essential for cancer progression and metastasis. We first measured the baseline cellular migration of male and female cells in a wound healing assay. At all timepoints measured, male GBM cells exhibited a significantly higher rate of cell migration over time compared to female cells **(Fig. 1D).** Based on these measurements, we chose the 12-hour timepoint as optimal to discriminate cellular migration from the potential confounding effect of cellular proliferation. We found treatment with SR18622 inhibited the migration of male cells, whereas female cells displayed a slight, non-significant increase in migration (**Fig 1E**), again demonstrating Egr1/Klf5 activity is necessary to maintain the observed sex difference in migration.

Finally, we sought to determine if the Egr1/Klf5 axis is required for sex differences in response to radiation exposure. Radiotherapy is the backbone of GBM treatment and male GBM patients and cells have been reported to exhibit more resistance to irradiation than females^4^, an observation which may explain the shorter progression-free survival in male compared to female GBM patients^2^. We treated both male and female GBM cells with either vehicle or increasing doses of SR18662 **(Supplemental Fig. 1)** in combination with 4Gy irradiation (4GyIR) and measured cell confluency over time using live cell imaging. SR18662 treatment (2µM) and radiation worked roughly additively on male (**Fig 1F, left panel**) cells (drug only led to ∼32% reduction, *p <0.0001*, radiation only led to ∼21% reduction, *p = 0.0011*, and the combination of the two led to ∼43% reduction in cell confluence) observed as early as 48hr post treatment. In contrast, while radiation reduced female cell proliferative capacity, drug treatment had no additional effect **(Fig. 1F, right panel and Supplemental Fig. 1B-E).** Thus, Egr1/Klf5 function appears to be a substantial component of the sex-biased proliferative response to radiation. Since radiotherapy has also been reported to trigger “radiation-induced cell migration” in GBM^31^, we next sought to examine the effect of drug treatment on cellular migration after radiation using the wound healing assay described previously. We found that male cells became more migratory after irradiation (26% wound closure after radiation versus 13% unirradiated, *p < 0.0001*), consistent with previous reports^32^, and this phenomenon was partly reversed by SR18662 (**Fig 1G)**. In contrast, female cells did not become significantly more migratory after IR treatment and drug treatment had no effect on this phenotype (**Fig 1G)**. Taken together, our results demonstrate Egr1/Klf5 function is necessary to maintain sex differences in proliferation, migration, and response to radiation in transformed astrocytes.

### SR18662 treatment has a sex-biased effect on cell invasion and clonogenic frequency in both murine transformed astrocytes and in human primary GBM cells

One hallmark of GBM is its aggressive nature; tumors metastasize quickly and have a high rate of occurrence, leading to poor outcomes. Two *ex vivo* cellular phenotypes, invasion and clonogenic frequency, are strongly correlated with poor outcomes, the former because it enables tumor cells to infiltrate surrounding tissues and lead to metastatic dissemination^33^, and the latter because clonogenic cells are indicative of a sub-population of self-renewing, tumor-recapitulating cells that are typically resistant to treatment^34^. We therefore sought to determine if Klf5/Egr1 activity is required for sex differences in these phenotypes. To quantify sex differences in invasion, we employed a trans-well invasion assay and analyzed male and female GBM cells, treated with either vehicle (DMSO) or drug (2µM SR18662). Trans-well invasion was significantly higher in male, compared to female GBM cells in our murine (mean number of invaded cells: 150 male; 115 female) model **(Fig. 2A).** Drug treatment (2µM SR18662) in male cells significantly reduced their invasive potential, rendering them similar to vehicle-treated female cells (**Fig. 2A**, mean number of invaded drug-treated male cells: 110), whereas drug treatment in female cells again had no significant effect on the phenotype (**Fig. 2A**, mean number of invaded drug-treated female cells: 120).

**Figure 2:**
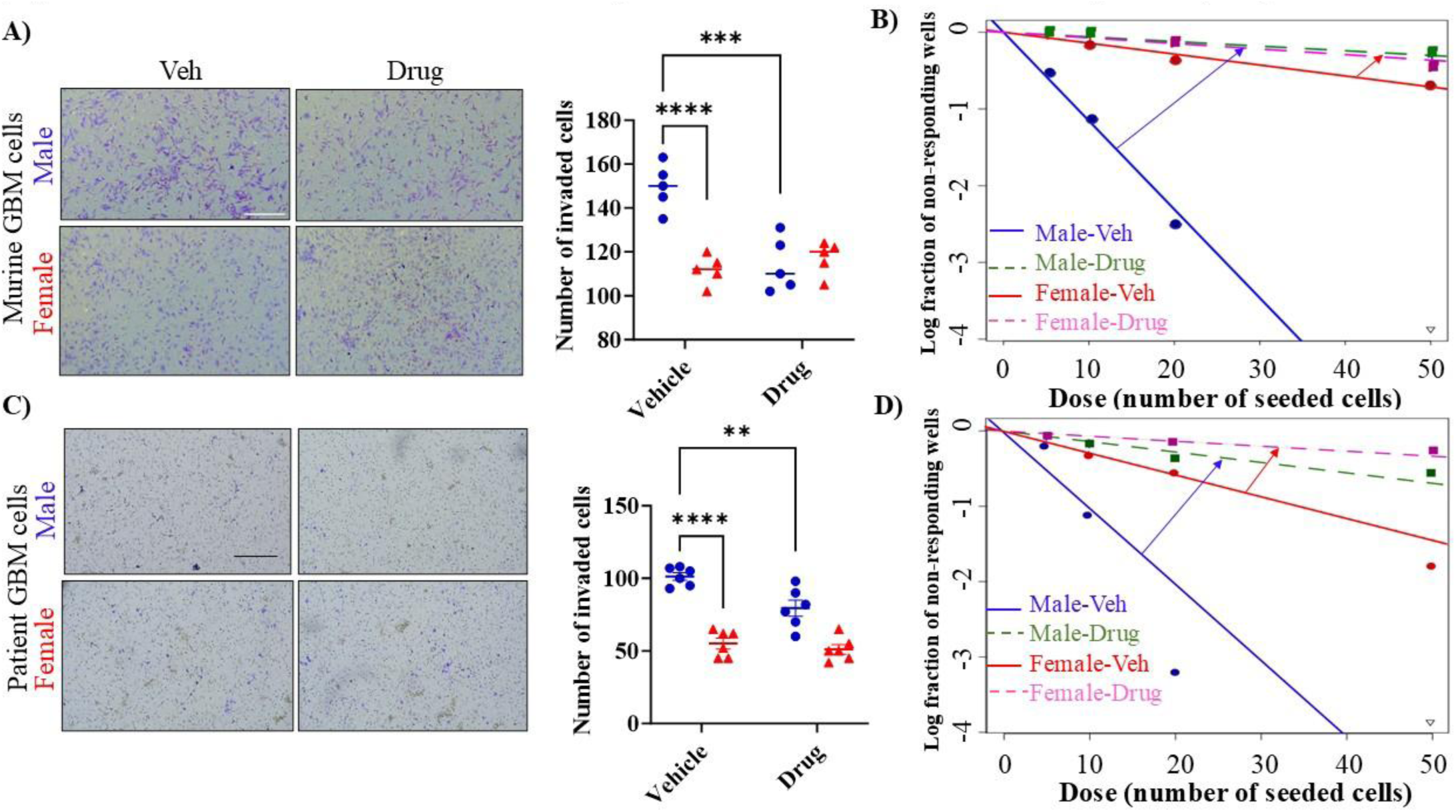
SR18662 treatment has a sex-biased effect on cell invasion and clonogenic frequency in both murine transformed astrocytes and in human primary GBM cells. **A)** Measure of trans-well invasion through Matrigel-coated chambers. Microscopic images of male and female murine *NF1-/- DNp53* (upper panel) and human (lower panel) GBM cells ± SR18662 (2µM) of crystal violet-stained cells after trans-well invasion assessed at 24hr. Scale bar = 200µm. Two-Way ANOVA on murine GBM cells: sex: *p < 0.0009*; males ± drug: *p = 0.0001*; females ± drug: *p > 0.999;* interaction: *p = 0.0002*. Two-Way ANOVA on human GBM patient cells: sex: *p < 0.0001*; males ± drug: *p = 0.005*; females ± drug: *p > 0.999;* interaction: *p = 0.035.* B) Measure of stem-cell clonogenic frequency using the extremely limited dilution assay (ELDA) in male (Veh: blue continuous line; drug: green dashed line) and female (Veh: red continuous line; drug: pink dashed line) murine *NF1-/- DNp53* (upper panel) and human (lower panel) GBM cells ± SR18662 (10µM) drug treatment. Sphere quantification conducted 14 days post seeding of single cells at 5, 10, 20 and 50 cells per well (24 x wells/density) and cultured in serum-free, stem-cell enriching media supplemented with hrEGF and hrFGF. Murine Males ± drug: *p <0.0001*; females ± drug: *p <0.05*; cell density seeded: *p < 0.0001*; interaction: *p < 0.0001*. Human patients GBM males ± drug: *p <0.0001*; females ± drug: *p <0.01*; cell density seeded: *p < 0.0001*; interaction: *p < 0.0001*. Experiments were conducted in triplicate biological repeats using 3x male and 3x female human patient GBM cells and analyzed using: http://bioinf.wehi.edu.au/software/elda/.

We next quantified the effect of SR18862 treatment on clonogenic stem cell frequency in male and female cells using the well-established Extreme Limiting Dilution Analysis (ELDA) assay. We used 10µM of SR18662 to ensure sufficient exposure for detecting subtle differences in stem cell frequency and drug sensitivity. The long-term (14-day) duration of the assay can reduce drug stability and efficacy over time, requiring a higher initial concentration to maintain consistent pressure on resistant stem-like cells. This approach enhances assay sensitivity and allows for a more accurate assessment of the drug’s impact on clonogenic potential. Male cells exhibited a significantly higher stem-cell clonogenic frequency than female cells as indicated by the steeper slope for male cells (**Fig 2B**, blue line) relative to female cells (**Fig 2B**, red line). Drug treatment (10µM SR18662, dashed lines) significantly reduced stem-cell clonogenic frequency in male cells (slope of green line versus red line, **Fig 2B**) but had little effect on female cells (pink line, **Fig 2B**). Taken together, our results again demonstrate that Klf5/Egr1 function is essential for maintaining sex difference in cellular invasion and clonogenicity.

Although our isogenic murine transformed astrocytes have been extensively validated as a model of GBM, it remains important to confirm findings in primary GBM lines. We therefore sought to determine whether Egr1/Klf5 downregulation reversed sex differences in human primary GBM cells. Human tumor cells behaved nearly identically to our murine model. Male cells were more invasive than female cells (**Fig 2C** and **Supplemental Fig 2 A and B**, quantification of sphere number and images), with a mean of 105 invaded cells for male lines versus a mean of 61 for female lines (*p < 0.0001*). SR18862 treatment (2µM) reduced invasion in male cells (mean: 78 cells, *p = 0.0053*), but not in female cells (mean: 56 cells, n.s.). The effects of SR18662 on clonogenic stem cell frequency in these human primary GBM cells also mirrored the results of the murine model: male GBM cells were more clonogenic than female GBM cells at baseline (**Fig 2D**, slope of blue line versus slope of red line), and after treatment with SR18662, male GBM cells became significantly less clonogenic (**Fig 2D**, slope of green line versus blue line) than did female cells (**Fig 2D**, slope of pink line versus red line). Thus, SR18662 has a strongly sex-biased effect on invasion and clonogenicity in our murine GBM model and in primary human GBM cells.

### Egr1, but not Klf5, knockout reverses sex differences in growth and migration and phenocopies SR18662 treatment

We next sought to determine if the ability of SR18662 to have this sex biased effect was due to on-target downregulation of the Egr1/Klf5 transcriptional pathway and not to an off-target interaction with some unknown protein or pathway. We used CRISPR to knockout (KO) Egr1 and Klf5, and polyclonal cell lines were used for phenotypic assays. We measured changes in tumorigenic phenotypes in male and female transformed astrocytes that were either vehicle or drug treated. We first measured changes in growth (**Supplemental Fig 3A**, lower panel of 12-day growth). Knockout of both Egr1 and Klf5 decreased growth in male cells, with Egr1 KO leading to a more significant effect than Klf5 KO (**Fig 3A**). Surprisingly, knockout of either Klf5 or Egr1 inhibited growth in male cells more potently than SR18862 treatment. To confirm that SR18862 acts through these pathways, we treated Klf5 KO and Egr1 KO cells with drug or vehicle (**Fig 3E, F** and **Supplemental Fig 3B-E**).We observed a significant additional reduction in growth in Klf5 KO cells after drug treatment (**Fig 3B**), but no additional reduction was observed Egr1 KO cells (**Fig 3C**), suggesting that SR18862’s inhibits growth in male transformed astrocytes via its ability to down-regulate Egr1. In female cells, KO of both Klf5 and Egr1 led to a small but significant increase in growth rate, whereas SR18862 treatment had no significant effect (**Fig 3D**), SR18862 treatment had no additional effect on the growth of Klf5 or Egr1 KO cells. The small differences in growth phenotype between SR18862 treated female cells and Klf5 KO or Egr1 KO cell are likely due to the fact that the CRISPR knockouts are have a bigger effect in growth than downregulation of these genes by SR18862.

**Figure 3:**
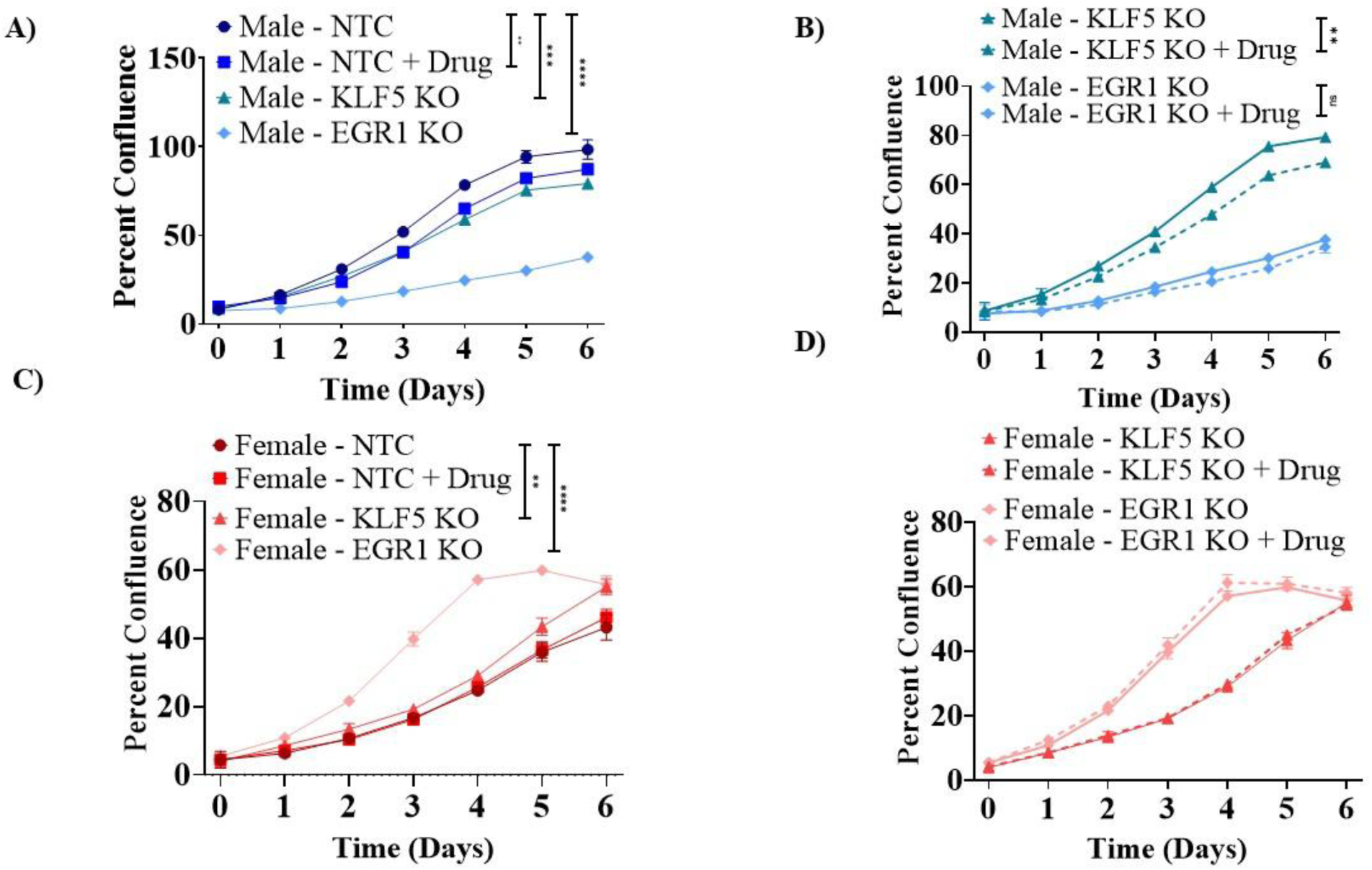
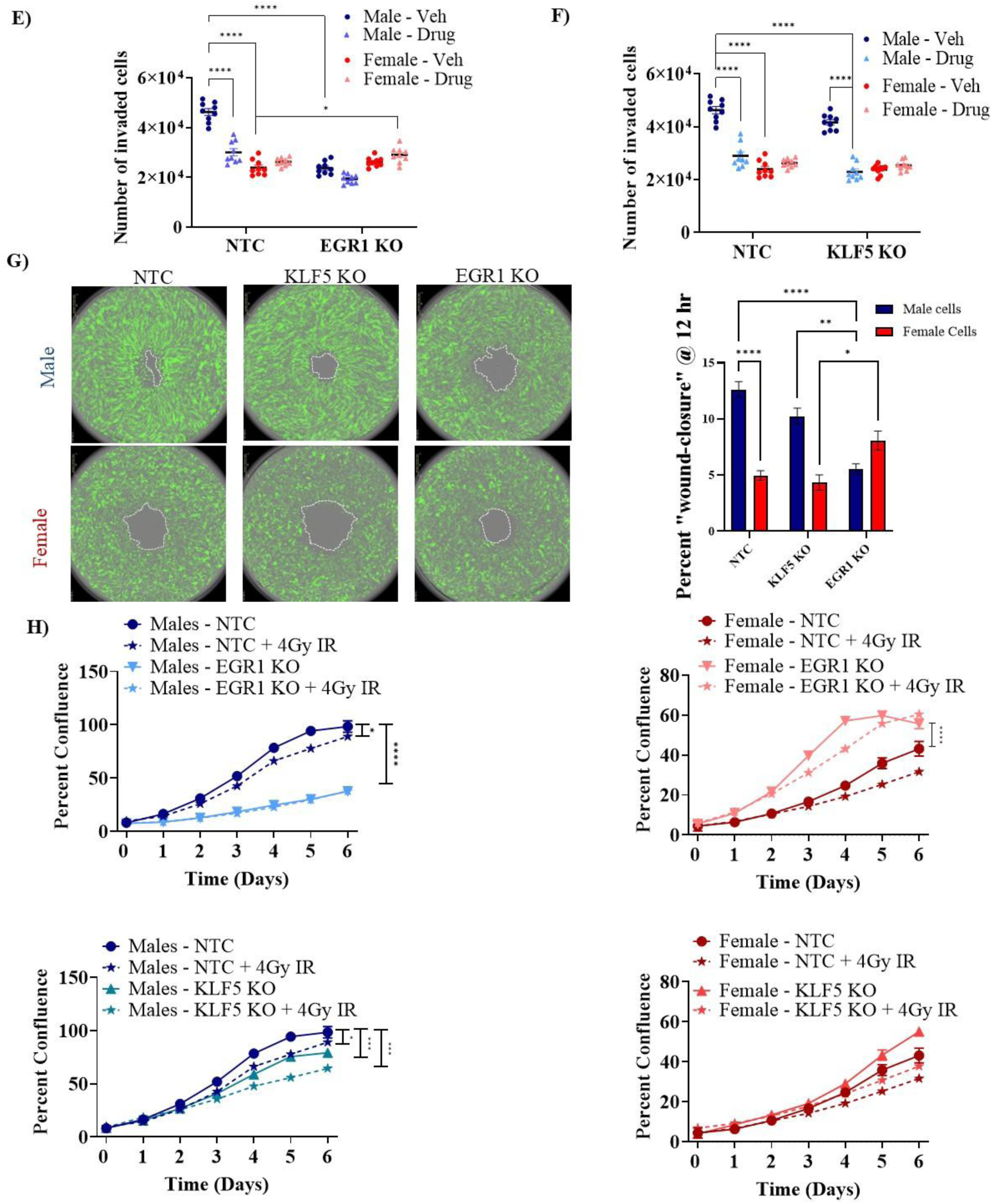
Egr1 is responsible for the sex-differences in murine GBM tumorigenic phenotypes. Time-course measure of cell confluency in male **(A and B)** and female **(C and D)** murine *NF1^-^/^-^ DNp53* cells ± Klf5 or Egr1CRISPR KO or drug (2µM SR18662) treatment. Three-way, repeated measures ANOVA at 48 hr: NTC versus Klf5 KO: *p = 0.39* (males) and *p < 0.05* (females), NTC versus Egr1 KO: *p < 0.0001*, time: *p < 0.0001*, sex: *p < 0.0001*, interaction: *P < 0.0001;* Drug: *p < 0.001* (males); *p > 0.999* (females); Klf5 KO ± drug: *p < 0.0001* (males); *p > 0.999* (females); Egr1 KO ± drug: *p > 0.999* (males and females). Quantification of trans-well invasion assay assessed @ 24 hr in male and female murine *NF1^-^/^-^ DNp53* cells ± Egr1 (**E) or** Klf5 **(F)** CRISPR KO and ± drug (2µM SR18662). Three-way ANOVA: Drug: *p = 0.008* (males only), sex: *p = 0.002*, interaction: *p = 0.0048*, NTC versus Klf5 KO: *p = 0.7005*, NTC versus Egr1 KO: *p = 0.001* (males only). **G)** IncuCyte live images and quantification of “wound-closure” in male and female murine *NF1^-^/^-^ DNp53* cells ± Klf5 or Egr1 CRISPR KO at 12 hr post “wound-induction”. Two-way ANOVA: sex: *p < 0.01*, Klf5 KO: *p > 0.999*, Egr1 KO: *p = 0.0001* (males), *p = 0.89* (females), interaction: *p < 0.0001*. **H)** Percent cell confluence in male (left panels) and female (right panels) *NF1^-^/^-^ DNp53* cells with Egr1 KO **(upper graphs)** or Klf5 KO **(lower graphs)** ± 4GY IR. IR: *p < 0.01*, Klf5 KO ± IR: *p < 0.001* (males); *p < 0.05* (females); Egr1 KO ± IR: *p < 0.0001* (females only). All experiments were conducted in triplicates and repeated at least twice. Error bars represent the mean of all biological repeats ± SEM.

We next investigated the effects of Klf5 and Egr1 KO on invasion. Klf5 KO did not significantly reduce invasive potential in male or female cells (**Fig 3F**). In contrast, Egr1 KO significantly reduced invasive potential in male (**Fig 3E**, ∼ 49% mean reduction in invaded cells, *p < 0.0001*) but not female transformed astrocytes (∼ 10% mean increase in invaded cells, n.s). The magnitude of the reduction with Egr1 KO is slightly higher than that achieved when unmodified male cells were treated with SR18862 (∼ 35% mean reduction, no significant difference), and treatment of male EGR1 K/D cells with SR18862 did not result in a further reduction of invasive potential. Together, these results provide evidence that SR18862 reduces the invasive potential of male cells by abrogating Egr1 activity in a manner that is independent of Klf5.

Egr1 and Klf5 KO had similar effects on cellular migration and response to radiation. Both knockouts reduced migration in male (**Fig 3G**, ∼50 and 20% mean reduction in invaded cells, *p<0.0001, p = n.s*) with Egr1 KO inhibiting cellular migration more potently than Klf5 KO in male cells, whereas in female cells, Egr1 tended to increase cell migration, albeit not significantly (**Fig 3G** and **Supplemental Fig 3F**). Finally, Egr1 KO significantly abrogated male cell response to radiation, as cell growth was reduced by > 50% (Fig 3H, upper left panel, *p < 0.0001*). In contrast, Egr1 KO female cells grew significantly faster than unmodified female cells after radiation treatment, such that the sex-bias after radiation was essentially reversed (**Fig 3H**). Klf5 knockout showed a similar trend, but the effect was more muted – male K/D cells grew more slowly than unmodified cells after radiation, whereas female KO cells grew faster after radiation (**Fig 3H**).

Taken together, our results suggest that SR18862’s ability to abrogate sex differences in growth, invasion, and migration are almost entirely due to downregulation of Egr1 and independent of Klf5 downregulation. However, SR18862 ability to abrogate sex differences in response to radiation appears to act through both Egr1 and Klf5.

### Egr1 knockdown leads to a decrease in proliferation and migration-related genes in male transformed astrocytes

Having established that SR18662 acts through Egr1 to inhibit male tumorigenic phenotypes such as proliferation, migration, invasion, and clonocgenicty, we next sought to understand the genomic profile in our murine model when we knocked down Egr1 (Egr1 KD). To do so, we knocked down Egr1 via a lentiviral system, utilizing the CRISPRi technology. Briefly, dCas9-KRAB transfected cells were selected under blastocidin until untransfected cells were almost 100% dead, before transfection with gRNA targeting Egr1 or a scramble gRNA as a control and selection with puromycin also in a similar manner. Three gRNAs targeting Egr1 were picked using the BROAD Institute’s CRISPick tool. Knockdown of Egr1 expression from the three guides tested was confirmed by qPCR analysis (**Supplemental Fig 4A**), whereas protein expression knockdown was confirmed with Western Blot (**Supplemental Fig 4B**). gRNA2 and 3 had the most knockdown for both males and females in both validation methods, and that is why they were used for further tests. RNA was collected from six replicates in a 6-well plate of each cell line targeting Egr1 or a scramble sequence (control), and libraries were prepared for bulk RNAseq (BRBseq^35^). There was a clear difference in 2-dimensional PCA plots in both sets of negative controls for both sexes (data not shown), with Negative control gRNA2 clustering with Egr1 KD cell lines, so only gRNA1 was used for downstream differential expression analyses. Additionally, gRNA2 and gRNA3 that target Egr1 were combined for differential expression (DE) analysis of Egr1 KD versus negative control in each sex. Unsurprisingly, males and females clustered separately in sample-sample distance heatmap (**Fig 4A**), and most of the most variance in the data was explained by sex, as both male and female cell lines, regardless of modification, clustered separated in PC1 (**Fig 4B**). DESeq2 was used to run differential expression (DE) analyses with a threshold of logFC of 0.5 and p-value of 0.05 used as cutoffs to determine significance. Thousands of genes were DE between male and female transformed astrocytes treated with Egr1 KD versus control cells of each respective sex.

**Figure 4:**
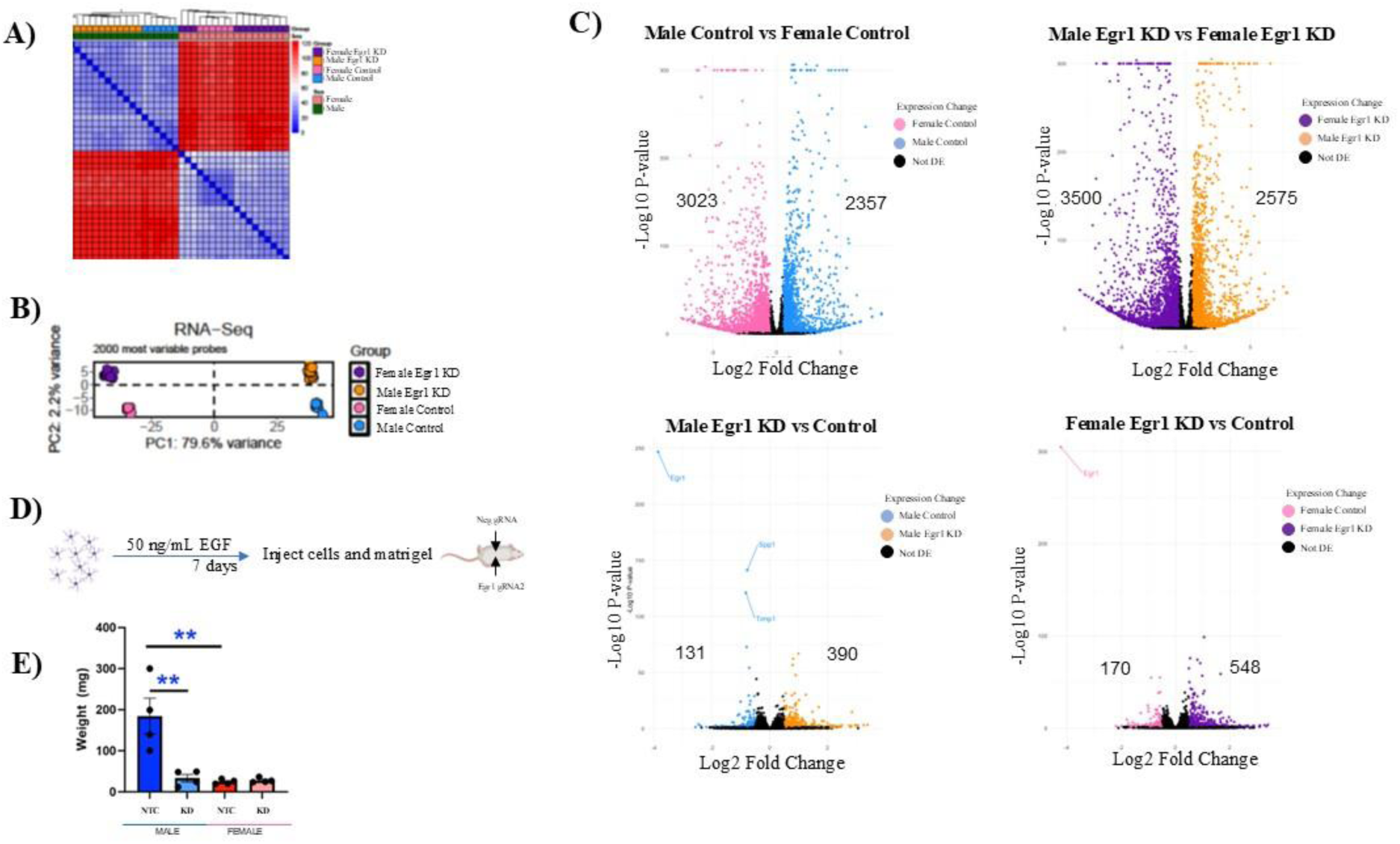
RNA-Seq analysis reveals sex-biased differential gene expression response to Egr1 KD. RNA-seq analysis of male and female murine *NF1-/- DNp53* GBM cells ± knockdown of Egr1. **A)** Heatmap of sample-sample similarity by gene expression. Males cluster together, shown in dark green to the left of the heatmap, whereas females cluster to the right shown in coral. Control male cell lines are shown in blue, and Egr1 KD in orange, whereas female control cell lines are shown in pink while the Egr1 KD are shown in purple. **B)** principal component analysis (PCA) of the 2000 most variable genes shows that 79.6% of variance in PC1 is explained by sex. PC2 shows differences within each sex upon knockdown of Egr1. **C)** Volcano plots illustrating log_2_ fold change on the x-axis and log_10_ p-values on the y-axis of DE genes in control male versus control female (upper, left panel), Egr1 knockdown males versus Egr1 KD females (upper, right panel), male Egr1 KD versus male control (lower, left panel) and female Egr1 KD versus female control (lower, right panel). D) Schematic of *in vivo* injections. Transformed astrocytes of each sex were treated with 50 ng/mL EGF for 7 days before injecting into the flanks of nude mice of the respective sex. One million male and two million female cells were infected with negative control gRNA2 or with Egr1 gRNA2, and each cell line mixed with matrigel was injected into the left or right flank of a nude mouse, respectively. E) Tumors were grown for 6 weeks, after which they were extracted from the mice and weighed. N=4/condition, **=p<0.005, Tukey’s multiple correction.

We observed large global transcriptional differences between negative control male and female cells (**Fig 4C, upper left plot**), with 3023 more significantly expressed in females, and 2357 genes more significantly expressed in males. Thousands of genes were also DE between Male and Female cell lines with Egr1 knockdown (**Fig 4C, upper right plot**), with 3500 genes more significantly expressed in females, and 2575 genes more significantly expressed in males. Knocking down this TF in male cells led to a significant decrease in expression of 131 genes, while 390 were significantly upregulated (**Fig 4C, lower left plot)**. In females, 170 genes were significantly downregulated, while 548 were significantly upregulated (**Fig 4C, lower right plot)**. More genes are upregulated in females upon Egr1 KD, showing that Egr1 inhibits more genes in females than in males.

To understand how the observed transcriptional changes contribute to the baseline phenotypic difference between male and female cells as well as how Egr1 knockdown reverses these differences, we performed a pathway analysis on the DEG sets discovered through our RNA-Seq analysis utilizing the enrichR package and the GO Biological Process 2023 database. Expected pathways were enriched in male cells compared to females, such as proliferation, DNA repair and epithelial to mesenchymal transition (EMT) **(Supplemental Table: 1),** whereas in females, significantly enriched pathways were migration, angiogenesis and proliferation (**Supplemental Table: 2).** Comparing male versus female cells after knocking down Egr1, the significantly upregulated genes that were enriched in males were again enriched for proliferation, DNA repair, and EMT (**Supplemental Table: 3**), whereas females were enriched for migration, angiogenesis and proliferation

(**Supplemental Table: 4**). Comparing Egr1 KD versus negative control in male cells, as expected based on our phenotypic results (**Fig 3**), genes that are significantly downregulated are involved in proliferation and migration (**Supplemental Table: 5**). Significantly upregulated genes in this comparison were enriched in the cholesterol pathway (**Supplemental Table: 6**), concluding that Egr1 inhibits the cholesterol pathway in male transformed astrocytes of our model. In females, significantly downregulated genes in the same type of comparison were enriched for epidermal growth factors and chemotaxis (**Supplemental Table: 7**) and an increase in cell migration and ECM (**Supplemental Table: 8**), again as expected in our phenotypic data (**Fig 3**). In conclusion, Egr1 upregulates pro-growth genes in males, while inhibiting pro-growth genes in females.

## Discussion

Glioblastoma is among the most aggressive cancers and is associated with dismal survival statistics. A long-standing mystery in the epidemiology of this disease is understanding why female patients have a lower incidence and better response to treatment than males. Here we provide evidence that Egr1 is a critical driver of this phenomenon.

Using an isogenic murine model of GBM, we show that the Egr1 and Klf5 motifs are enriched in stretch and regular enhancers, respectively. As Egr1 directly regulates Klf5, and our multivariate Cox analysis from human patients with GBM data from TCGA revealed that EGR1 expression levels affect survival in GBM patients, we naturally decided to study these two TFs together. We show here that the Egr1/Klf5 inhibitor SR18662 leads to a sex-biased phenotype across multiple tumorigenic phenotypes: cell growth, clonogenicity, migration, invasion and response to radiation. In all instances, SR18662 abrogates the phenotype in male cells, but has little effect in female cells. These results were extended to human primary GBM cells, demonstrating that the effects of Klf5/Egr1 downregulation are not limited to our murine model and the phenotypic consequences are large enough that they are not masked by the genetic and epigenetic heterogeneity inherent in patient-derived lines. To determine whether Egr1 or Klf5 is the crucial player in the observed effects, we knocked out each TF individually and found that inhibiting Egr1 best phenocopied the observed sex differences in cancer phenotypes after SR18662 treatment. Importantly, in male cells, the addition of SR18662, decreased confluence in Klf5 KO cells, while it had no further effect in Egr1 KO cells. In contrast, the addition of the drug had no effect on female cells with or without Egr1 or Klf5 knockout. As such, we conclude that sex differences in the key tumorigenic phenotypes of growth, clonogenicity, invasion, and migration are largely mediated by Egr1 in a manner that is independent of Klf5 downregulation. The only phenotype where Klf5 may play a significant role is response to radiation, but even then, our results suggest that Egr1 plays an additional role in maintaining sex differences beyond Klf5 regulation.

Bulk RNAseq of Egr1 knockdown cell lines of our murine model showed expected changes in gene expression. Male cells with Egr1 KD showed downregulation for pro-tumorigenic genes related to proliferation and migration, whereas those downregulated in female Egr1 KD cell lines were enriched in pro-tumorigenic pathways like migration and ECM. Interestingly, Egr1 plays different roles in different contexts, as it can promote or inhibit tumorigenesis based on the cell type it is expressed in. Our results here indicate that a sex component might need to be added to these cancers that display sex differences. Further studies need to be conducted to determine downstream targets of Egr1 that play a role in these sex differences in our murine GBM model as well as in human cell lines with methods such as Calling Cards^36^ and CRISPRi downregulation.

## Conclusions

We conclude that Egr1 plays a major role in maintaining sex difference across multiple tumorigenic phenotypes in GBM. Downregulation of Egr1 reduces growth, clonogenicity, migration, invasion, and response to radiation in male GBM cells. In contrast, in female cells, these phenotypes are unaffected, or in some instances, slightly enhanced. This oppositional role of Egr1 sheds light on the possible mechanisms by which sex-differences in tumorigenic phenotypes arise and warrants further investigation to deliver more efficacious cures to patients with GBM.

## Materials and Methods

### Patient Survival Analysis

Human GBM patient gene expression data was downloaded from glioVis. Multivariate Cox proportional hazard model with the following covariates (EGR1, sex, age, MGMT methylation status, and recurrence) was performed on TCGA GBM cohort data from glioVis. Analyses were conducted of All patients with IDH1 WT tumors combined (n = 271), and in only male (n = 157) or female (n = 114) patients separately.

### Cells and Media

Male and female GBM (*Nf1^−^/^−^; DNp53*) astrocytes were generated as previously reported^6^ and cells were grown in DMEM/F12 media supplemented with 10% FBS and 1% penicillin-streptomycin. Chemical inhibition of KLF5/EGR1 was achieved using the drugs labeled as “KLF5 inhibitors” CID 5051293 (Sigma-Aldrich Cat # 422625) or SR18662 (SelleckChem. Cat # Catalog No. S8900). While these drugs are labelled as “KLF5 inhibitors”, their specific mechanism of action in via down-regulation of EGR1 protein expression that subsequently leads to reduced transcription of KLF5. Drugs were reconstituted in DMSO to 131 mM or 25 mM, respectively, and used at the indicated doses.

### Cell Viability using trypan blue exclusion assay

Male and female *(Nf1^–^/^–^ DNp53)* murine GBM cells were grown in 6-well plates (50,000 cells per well). At the designated time-points, cells were washed with cold PBS, trypsinized for 5 minutes to detach cells from the culture flask and then neutralized with full-serum media. The cell/trypsin/media mixture was centrifuged at 300 x g for 5 minutes and supernatant discarded after which cells were resuspended in full-serum ready for quantification. Cell viability was measured every 24 hours by Trypan Blue exclusion (final concentration 0.1%) using a Countess 3 FL Automated Cell Counter (Invitrogen; Cat # A49893) with triplicate wells per time point per sex up to 96 hr post seeding.

### Cell confluency assays

Male and female *(Nf1^–^/^–^ DNp53)* murine GBM cells ± KLF5 or EGR1 CRISPR K/D or ± SR18662 at the indicated dosages were seeded in full-serum media at a density of 1000 cells/well of a 96-well plate with 6 replicates per condition. Cell confluency was assessed over time (0-6 days) using the IncuCyte ZOOM live cell imaging platform (Sartorius). Continuous live videos and phase-contrast (and green fluorescent) images of the whole well were acquired for qualitative representation of cell confluency between the treatment groups every 12 hr for 6 days and cell confluency analyzed using IncuCyte ZOOM analysis software. Percent cell confluency, as a measure of longitudinal cell growth, was graphed using GraphPad Prism and analyzed with repeated measures ANOVA.

### Cell migration assays

Male and female *(Nf1^–^/^–^ DNp53)* murine GBM cells were seeded in full-serum media at a density of 20,000 cells/well in a 96-well plate in which a circular plug was inserted in the middle of each well. Cells were allowed to seed around the periphery of the plug (to mimic a potential “wound” in the middle of the 96-well plate) for 6 hr before removing the plug. Cells were placed in the incubator housing the IncuCyte (Sartorius) platform that acquired images of the “wound” area designated by the circular, cell-free region where the plug had been inserted, every 6 hr for 3 days. “Wound-healing” was measured by the cells’ ability to migrate into and close the “wound” over the 3-day period. The “wound” area was measured using AxioVision software (Ziess) at 0 time-point and then at 6, 12, 24, 36, 48 and 60 hr. To calculate the percentage of “wound-closure”, the following formula was used: % of wound closure = [(wound area at 0 h – wound area after X h)/wound area at 0 h] x 100.

### Cell invasion assays

Male and female *(Nf1^–^/^–^ DNp53)* murine GBM cells or human primary GBM cells were grown in full-serum media in T75 flasks until approximately 70-80% confluency and then washed 2x with serum-free media and cultured over night with serum-free media to starve the cells before induction of cell invasion. The next day cells were treated with either vehicle (DMSO) or drug (SR18662) for 1 hr, then collected using trypsin and counted on a Countess automated cell counter. One hundred thousand cells were seeded into the top of a Matrigel coated invasion chamber in 500µL serum-free media and full-serum media added to the wells on the underside of the invasion chamber. Cell invasion through the Matrigel coated chamber towards the full-serum on the underside of the chamber was calculated after 24 hr. To quantitate cell invasion, the un-invaded cells were removed from the upper side of the chamber using cotton tips and the invaded cells that were adhered to the bottom side of the invasion chamber were fixed with ethanol and stained with crystal violet, then imaged using InCell (GE InCell Analyzer 6500, Cellomics Inc) live cell imaging platform and counted using ImageJ^37^.

### Stem-cell clonogenic frequency assay

Male and female *(Nf1^–^/^–^ DNp53)* murine GBM cells or human primary GBM cells were collected and resuspended in stem-cell, serum-free media supplemented with hrEGR and hrFGF. Cells were seeded at very limited dilution (100, 50, 20, 5 cells per well) in stem-cell, serum-free media supplemented with rhEGF and rhFGF and heparin into 96-well ultralow cluster plates (Corning), 24 wells per concentration using the CytoFlex SRT cell sorter (Beckman). Sphere formation was measured 14 days after plating. Clonogenic stem-like cell frequency was analyzed using the Extreme Limiting Dilution Analysis (http://bioinf.wehi.edu.au/software/elda/).

### Cell Irradiation

For cellular irradiation experiments, cells were irradiated using an RS 2000 X-ray irradiator (Rad Source Technologies). Radiation was delivered at a dose rate of ∼2, 4, 8 Gy/min with 160 kVp X-rays. Unirradiated control cells were transported to the irradiator together with the irradiated cells and sat on the bench for the same length of time as the cells that were in the irradiator.

### CRISPR Knockout

To knockout KLF5 or EGR1, we employed CRISPR ribonucleoproteins (RNPs) using the CRISPRevolution sgRNA EZ Kit (1.5 nmol) to target either protein according to the manufacturer’s protocol (Synthego, Inc.). For KLF5, three independent RNPs (KLF5gRNA #1 CUCAUGGUCAGCACCCGCGU, KLF5gRNA #2 AGCACCCGCGUGGGCAUGGA and KLF5gRNA #3 GGUCAGCACCCGCGUGGGCA) were used, and for EGR1 knock-down, two independent RNPs (Egr1gRNA #1 CAUCAAUUGCAUCUCGGCCU and Egr1gRNA#2 CGGAGACAUCAAUUGCAUCU) were used. Scrambled non-targeting gRNA was used as control (NTC). Briefly, Male and female murine GBM cells were cultured in T-25 flasks with full-serum until ∼ 70% confluency was reached. CRISPR RNP (2x sgRNA for EGR1, 3x sgRNA for KLF5 and 1x sgRNA for scrambled, non-targeting control) complexes (1.3:1 sgRNA to Cas9 ratio) were prepared by mixing the sgRNA, Cas9 and Lipofectamine Cas9 Plus reagent (Synthego, Inc.) with Opti-MEM reduced serum medium and set aside for 5-10 minutes. The transfection solution was prepared by mixing Lipofectamine CRISPRMAX reagent with Opti-MEM reduced serum medium and set aside for 5 minutes. The two solutions were mixed together with gentle pipetting and set aside for 10 minutes. Cells were trypsinized collected, counted and ∼ 100K viable cells plated per reaction in a 12-well plate in a “reverse transfection” reaction with the gRNA-transfection solution to achieve KLF5 or EGR1 knockout for 24 hr before cells were washed and transfection reagent replaced with full-serum media. Cell pellets were collected 72 hr post transfection and stored in RNA Protect for future RNA extraction for qRT-PCR quantification of mRNA or RIPA cell lysis buffer for Western blot analysis of protein expression.

### CRISPRi knockdown

Before every transfection, transformed mouse astrocytes were seeded using regular growth media and grown overnight. First, cells were transfected with dCas9 plasmid for 24 hrs, after which, media was changed, and cells were left to rest overnight. The next day, cells were split and selection with blastocidin was started at the same time. When the negative controls had died up to 95%, cells were split and let to rest for 1 passage before infecting with gRNA lentivirus, with media without selection antibiotic. After infection with gRNA lentivirus, the selection media contained blasticidin as well as puromycin. After selection was done, cells were used for bulk RNA expression experiments and in vivo injections in nude mice.gRNAs were designed using the BROAD Institute’s CRISPick tool (RRID:SCR_025148). Three indeependent sets of gRNAs were used to knockdown Egr1 (gRNA1: GGTTGGCCGGGTTACATGCG, gRNA2: GGCAGGGGCCGATCTTGCGG, gRNA3: GCGGCGGCGGCGAATCGCGG). Knockdown was confirmed with qPCR using the following primers: Egr1_mouse_qPCR_FP: cagcgccttcaatcctcaag and Egr1_mouse_qPCR_RP: gagaagcggccagtataggt.

### Western Blots analysis

Total cell protein lysates were collected with 0.25% trypsin and were washed with cold PBS. Cells were lysed with RIPA cell lysis buffer composed of 1% NP-40, 50 mM Tris-HCl pH 8.0, 150 mM NaCl, 5 mM EDTA, 0.5% sodium deoxycholate, 0.1% SDS together with Halt™ Protease and Phosphatase Inhibitor Cocktail (100X) (Thermo Fisher Scientific, 78440). Samples were prepared using NuPAGE™ Sample Reducing Agent (Invitrogen, NP0004) and urea loading buffer (5x: 8M urea, 10% w/v SDS, 10 mM 2-mercaptoethanol, 20% v/v glycerol, 0.2M Tris-HCl pH 6.8, 0.05% w/v bromophenol blue) and were loaded on a NuPAGE™ Bis-Tris protein. The following antibodies were used: Egr1 (44D5) Rabbit mAb #4154 and Beta-actin (CST β-actin (1:5000, 8457, RRID:AB_10950489) and secondary antibodies anti-rabbit (1:2500, 7074, RRID:AB_2099233 from Cell Signaling Technologies).

### Quantitative RT-PCR

RNA was isolated from cells using the RNeasy Plus Mini Kit (Qiagen, 74136) according to the manufacturer instructionsRNA concentration was quantified using a NanoDrop 1000 spectrophotometer (Thermo Scientific), and 120 ng of RNA was used to synthesize cDNA using Invitrogen’s SuperScript™ VILO™ cDNA Synthesis Kit (ThermoFisher Scientific, 11754250). Quantitative RT-PCR was performed using PowerUp™ SYBR™ Green Master Mix (ThermoFisher Scientific, A25741) on the QuantStudio 3 Real-Time PCR (ThermoFisher Scientific).

### RNA-Seq library construction

RNA-Sequencing to determine differential gene expression after Egr1 KD was conducted by extracting RNA from Egr1 KD using the Qiagen RNAeasy Plus Micro kit (Qiagen, Cat # 74034) according to the manufacturer’s protocol. Briefly, poly-adenylated transcripts were annealed to molecular barcodes (IDT, Inc.) containing a 16-nucleotide sequence unique to each 0.2mL reaction vessel. Barcoded transcripts were reverse-transcribed to first-strand cDNA. The hybrid transcript:cDNA molecules from every sample was pooled for subsequent cDNA amplification. 2ng of amplified cDNA was used for library construction with Illumina Nextera XT protocol (https://genomebiology.biomedcentral.com/articles/10.1186/s13059-019-1671-x). Multiplexed 3’mRNA-seq libraries were made using the Alithea BRB-seq kit (Alithea Genomics, Cat #10813). XX ng of RNA was used as input for every sample. The final sequencing library was made by tagmenting 20 ng of cDNA input with 3 uL of the TE reagent. Adapters and polyA ends were removed using CutAdapt (2.10). Reads were aligned to the mouse genome (GRCm38), demultiplexed, and UMI de-duplicated using the STARsolo mode of STAR (2.7.6a).

### Bulk RNA-sequencing analysis

Genes were filtered by requiring a minimum Counts Per Million (CPM) threshhold of 1to remove all zero-count data. The top 2000 most variable genes were used for principal component analysis (PCA), and Euclidian Distance was used to calculate sample-sample distance in the heatmap. Differential expression (DE) analysis was performed using DESeq2 (version 1.42.1). A threshold of logFC of 0.5 and p-value of 0.05 was used to determine significance in DE genes. Gene enrichment analysis was run using the enrichR package in R (version 3.4) was used to run gene enrichment, using the GO Biological Process 2023 database.

### Xenograft experiments

One million male and 2 million female control or Egr1 knockdown cell lines from our GBM mouse astrocyte model were injected into the left and right flanks of recipient mice of the same sex, respectively. Four mice were used for each injection, and tumors were left to grow for 6 weeks, after which tumor size was determined.

### Statistical analysis

Experiments were run in triplicate and repeated at least twice. Analysis of variance (ANOVA) with Bonferroni’s correction for multiple comparisons test was used to establish significance (alpha at *p<0.05)*. One-way ANOVA was used to examine statistical significance in cell proliferation, viability, migration and invasion assays between male and female murine and human GBM cells. A two-way ANOVA was used to examine statistical significance in phenotypic behavior or protein expression between vehicle- or drug-treated male and female cells and a three-way ANOVA used when drug or IR were combined with or without EGR1 or KLF5 knockdown. Error-bars represent mean ± SEM of multiple experiments.

## Data availability

The data generated in this study is available upon request.

## Acknowledgements

The authors wish to acknowledge the personnel of the Rubin and Mitra labs for their support throughout this work. The authors would also like to thank the personnel at the DNA Sequencing Innovation and High Throughput Computing Facility at The Edison Family Center for Genome Sciences and Systems Biology of Washington University in St. Louis for their sequencing and computational expertise.

## Author contributions

Conceptualization, TA, LLl, JR, and RM; methodology, TA and LLl; software, RM; validation, TA, LLl, JPW, LY, NMW, LT, BM; formal analysis, TA, LLl and JL; investigation, TA and LLl; resources, JR and RM; data curation, TA and LLl; writing LLl, TA, JR, RM; supervision, JR and RM; project administration, JR and RM; funding acquisition, JR and RM. All authors have read and agreed to the published version of the manuscript.

---

**Supplemental Figure 1:**
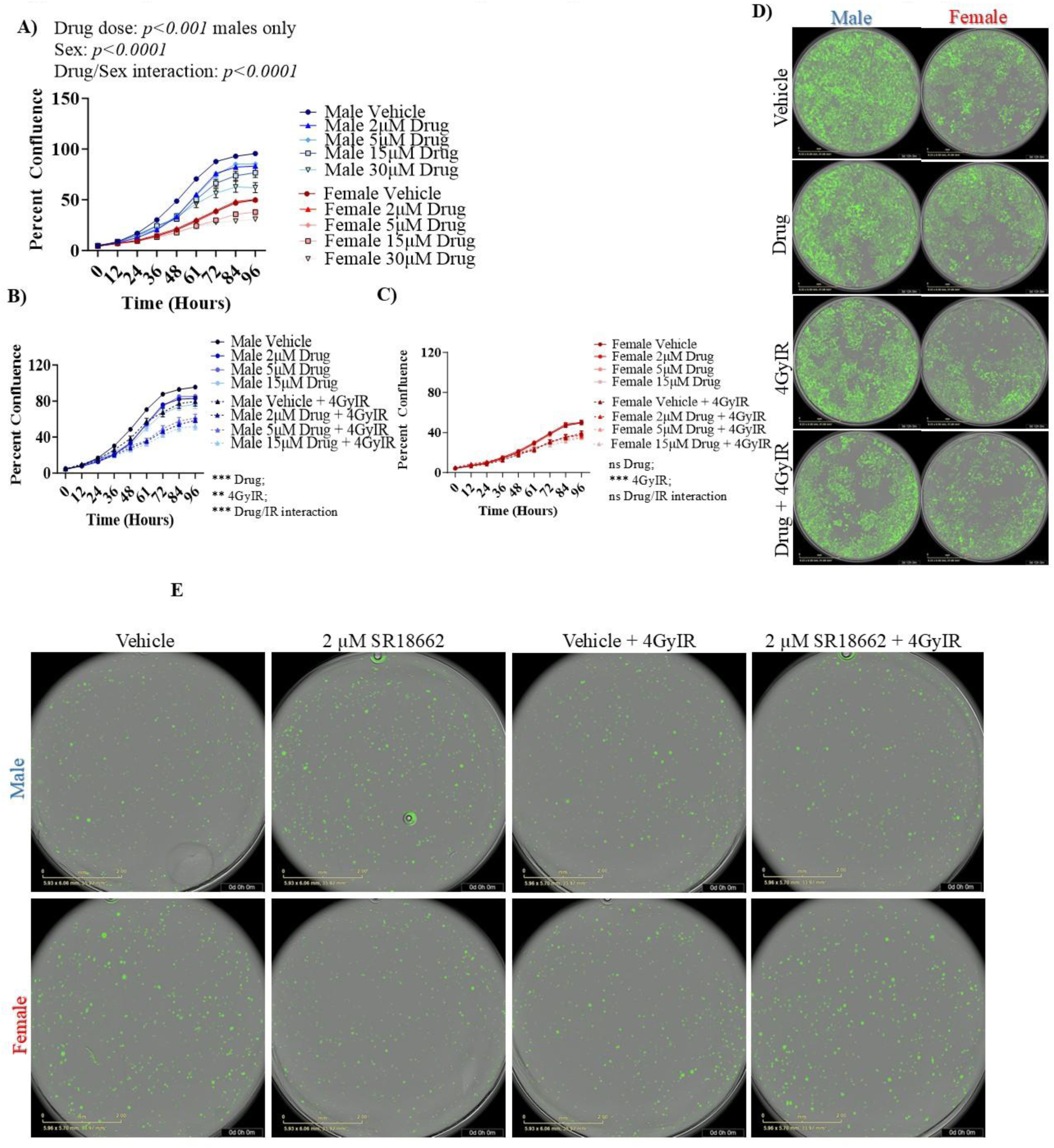
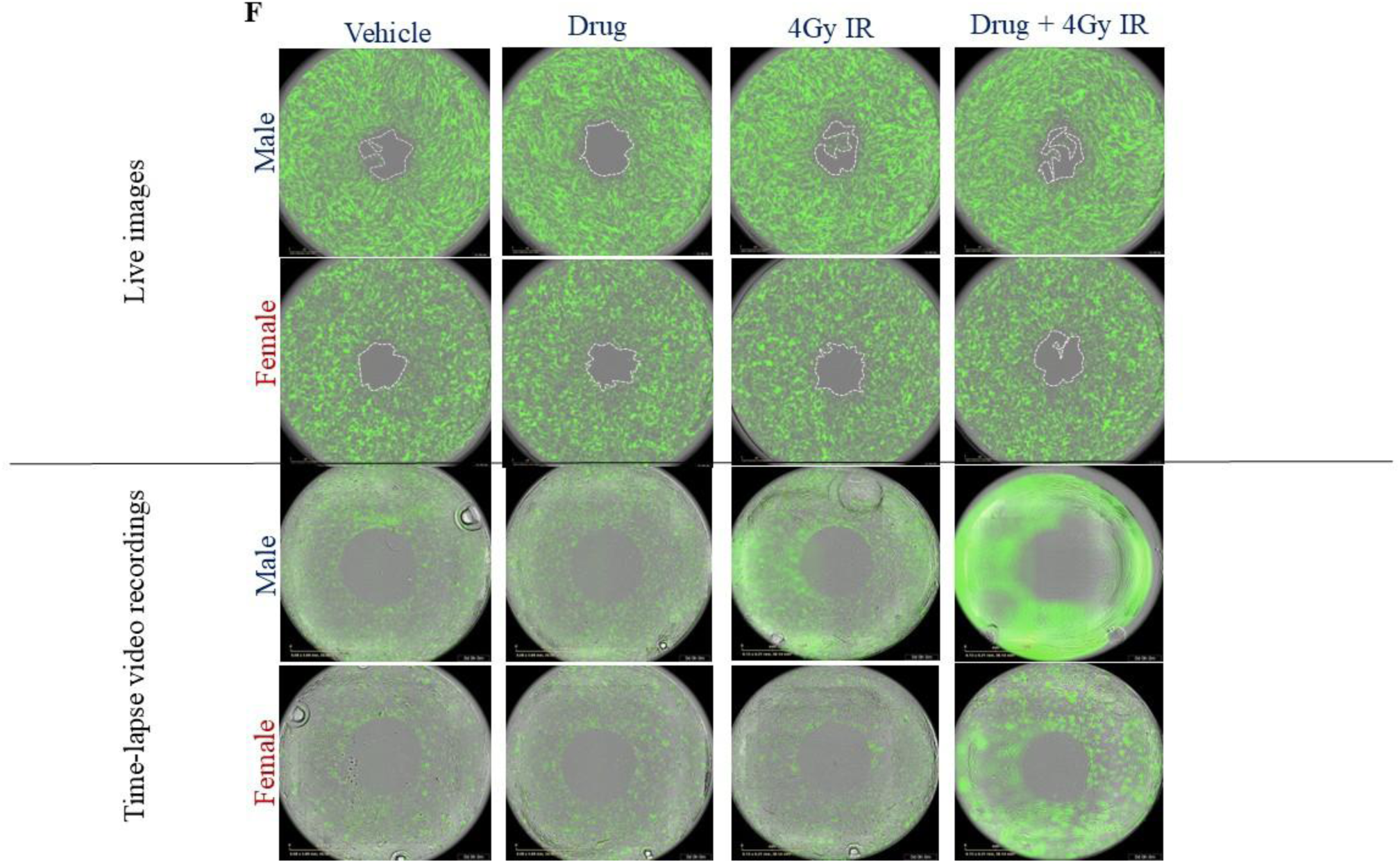
Tumor cell sex determines dose-dependent response and radio-sensitization to SR18662 treatment. **A)** Cell confluency measured over time (96 hr) in male (blue-shaded lines) and female (red-shaded lines) murine *NF1^-^/^-^ DNp53* treated with increasing doses of SR18662 (2-30µM). Cell confluency measured over time (96 hr) in male **(B)** and female **(C)** murine *NF1^-^/^-^ DNp53* treated with SR18662 (2, 5 and 15µM) plus/minus 4GyIR. Live images taken of the 2µM SR18662 dose at 48hr post IR **(D)** and time-lapse video recordings over 5 days **(E)** in male and female murine *NF1^-^/^-^ DNp53* cells. **F)** Live images and time-lapse video recordings of “wound-healing” in male and female murine *NF1^-^/^-^ DNp53* cells ± drug, ± 4GyIR. *p<0.05; **p<0.01; ***p<0.001; ****p <0.0001.

**Supplemental Figure 2:**
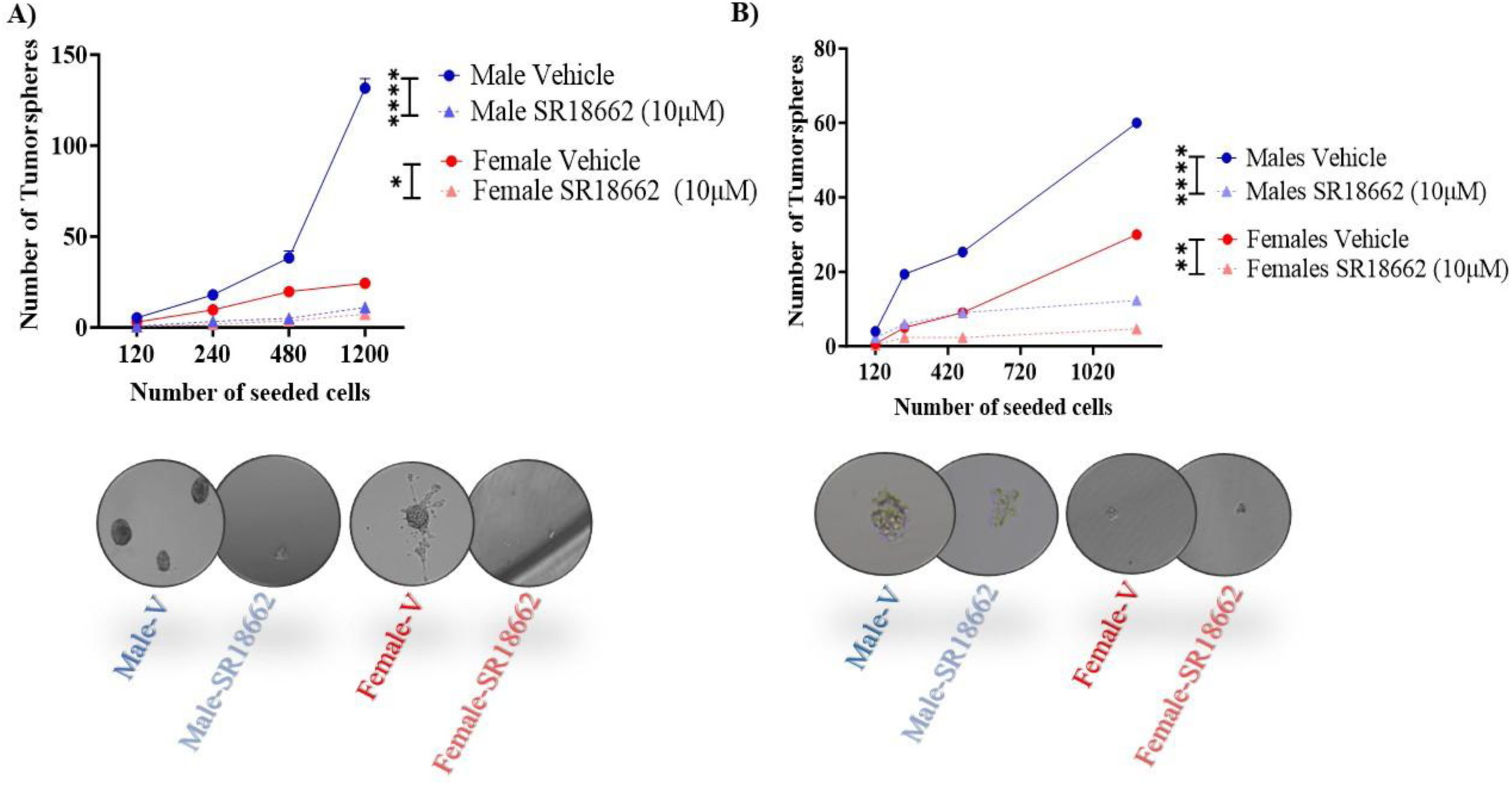
SR18662 treatment induces a sex-biased response on cell invasion and stem-cell clonogenic frequency in GBM. Graphical representation of the number of spheres formed per number of cells seeded in murine *NF1^-^/^-^ DNp53* **(A)** and human **(B)** GBM cells and representative live images of tumorspheres (images below respective graphs). *p<0.05; **p<0.01; ****p <0.0001.

**Supplemental Figure 3:**
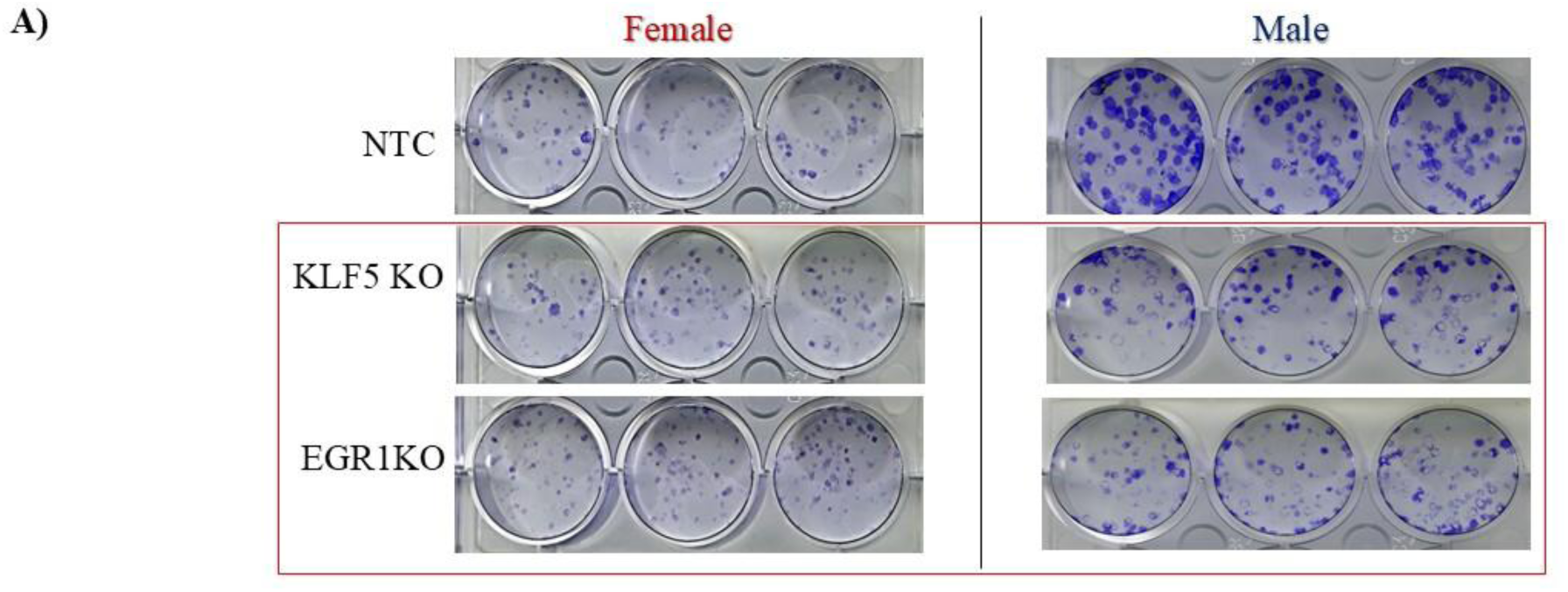
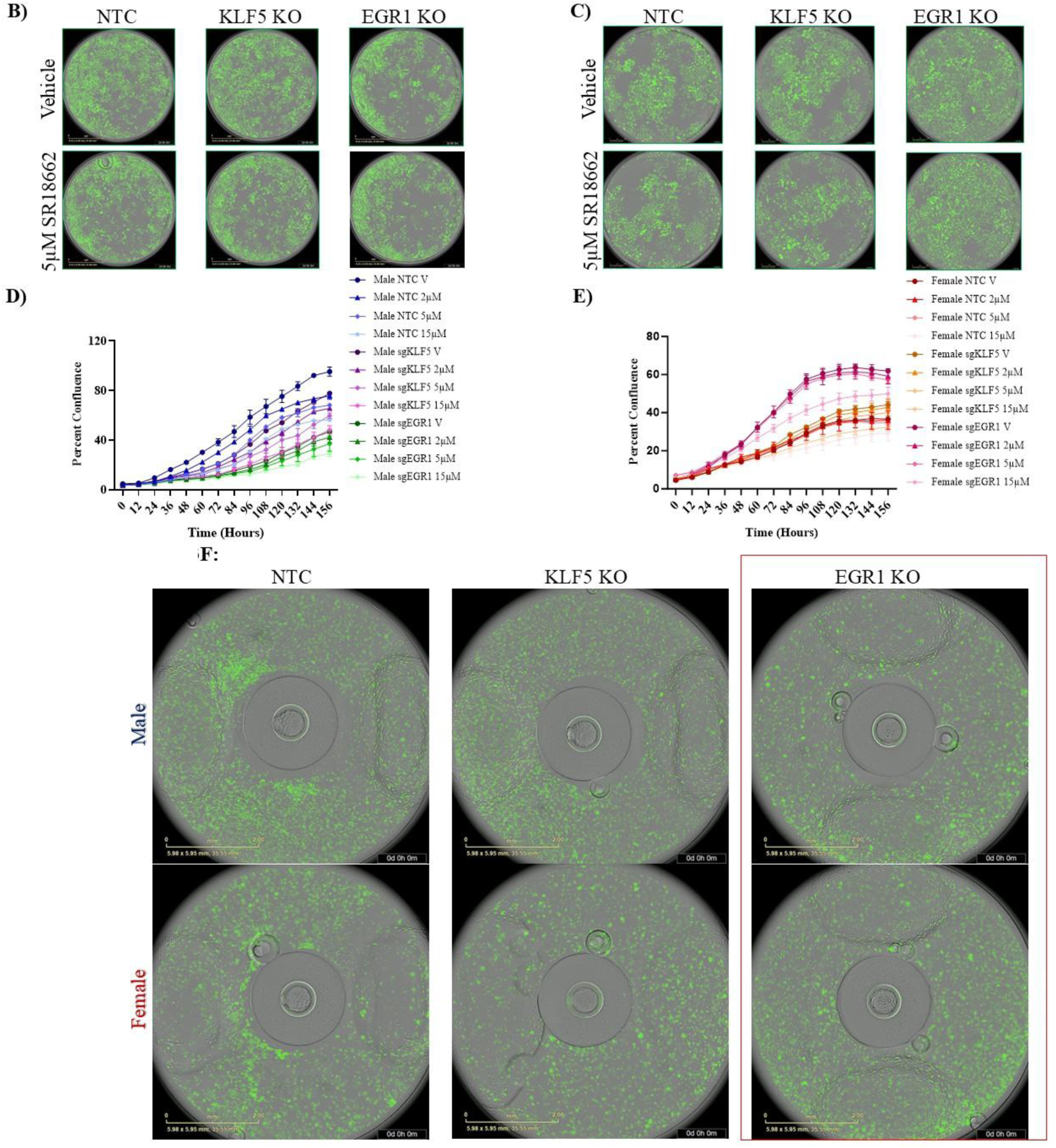
Egr1 is responsible for the sex-differences in murine GBM tumorigenic phenotypes. **A)** Images of long-term cell growth (12 days) in male and female murine *NF1^-^/^-^ DNp53* GBM cells after KLF5 or EGR1 K/D compared to NTC cells. Live cell images **(B)** and graphical representation of male **(D)** and female **(C and E)** murine GBM cells with NTC, Klf5 or Egr1 CRISPR KO, treated with either vehicle or increasing doses of SR18662. Time-lapse video recording of “wound-closure” **(F)** in murine male (upper panel) and female (lower panel) *NF1^-^/^-^ DNp53* GBM cells after Klf5 or Egr1 KO compared to NTC.

**Supplemental Figure 4:**
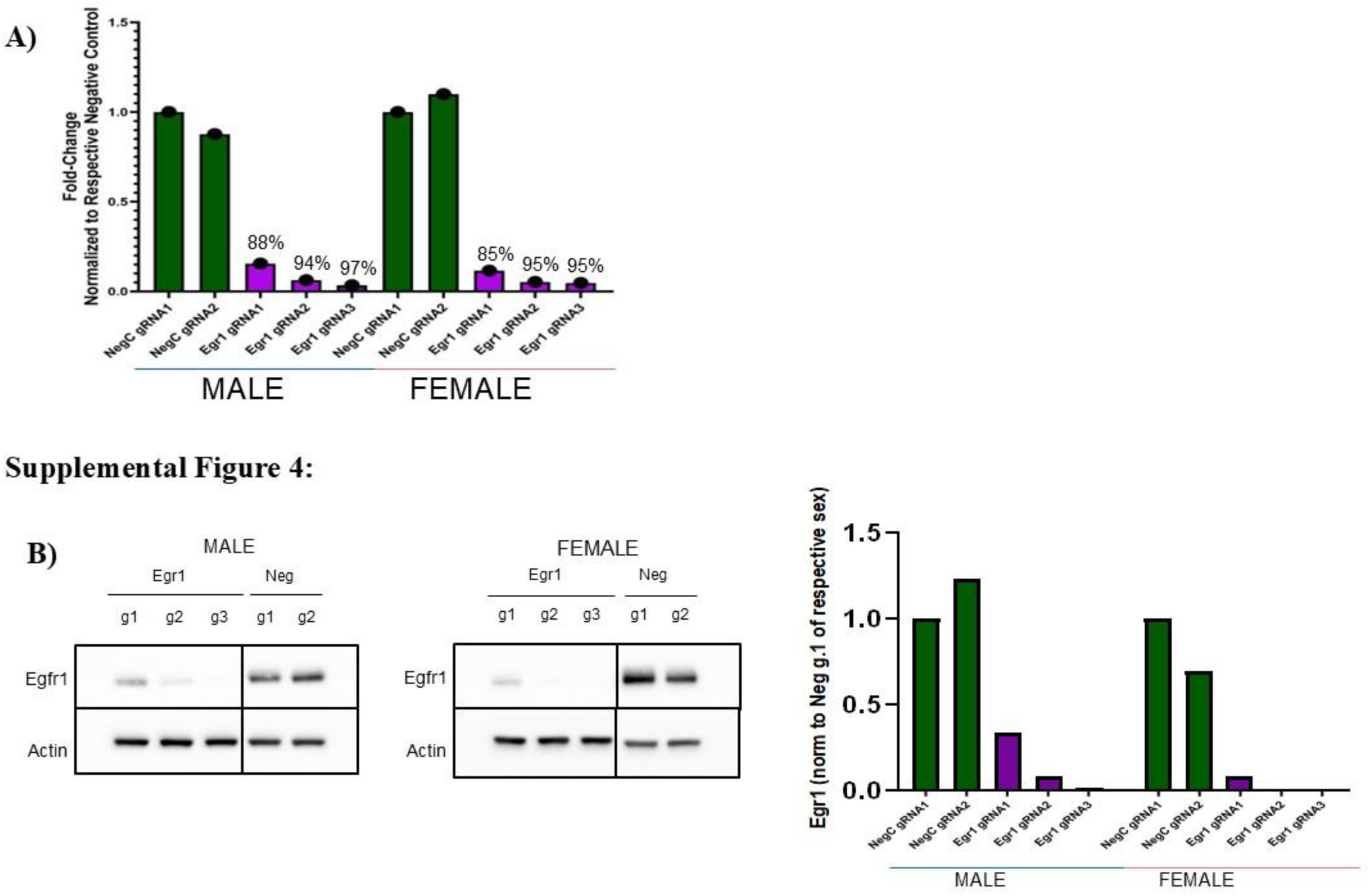
RNA-Seq analysis reveals sex-biased differential gene expression response to Egr1 KD. **A)** qPCR results to show Egr1 KD using three gRNAs targeting Egr1 and 2 scramble gRNAs as negative control in male and female murine model of GBM. **B**) Western blots (left figure) showing protein expression of Egr1 KD and control cell lines in male and female murine model of GBM and quantification (right panel).

